# Dietary regulation of silent synapses in the dorsolateral striatum

**DOI:** 10.1101/2024.03.24.586457

**Authors:** Allison M. Meyers, Federico G. Gnazzo, Eddy D. Barrera, Tikva Nabatian, Larry Chan, Jeff A. Beeler

**Author notes:** **Competing interests:** The authors have declared no competing interests.

## Abstract

Obesity and drugs of abuse share overlapping neural circuits and behaviors. Silent synapses are transient synapses that are important for remodeling brain circuits. They are prevalent during early development but largely disappear by adulthood. Drugs of abuse increase silent synapses during adulthood and may facilitate reorganizing brain circuits around drug-related experience, facilitating addiction and contributing to relapse during treatment and abstinence. Whether obesity causes alterations in the expression of silent synapses in a manner similar to drugs of abuse has not been examined. Using a dietary-induced obesity paradigm, mice that chronically consumed high fat diet (HFD) exhibited increased silent synapses in both direct and indirect pathway medium spiny neurons in the dorsolateral striatum. Both the time of onset of increased silent synapses and their normalization upon discontinuation of HFD occurs on an extended time scale compared to drugs of abuse. These data demonstrate that chronic consumption of HFD, like drugs of abuse, can alter mechanisms of circuit plasticity likely facilitating neural reorganization analogous to drugs of abuse.

## INTRODUCTION

Obesity has increased in Western society in recent decades with a prevalence of ~42% in adults (Flegal et al., 2016). Most individuals who attempt to diet tend to fail in the long term, returning to their original weight or, often, rebounding to a higher weight (Dulloo et al., 1997; Speakman et al., 2011). This failure to maintain diet and weight loss may be caused by the compulsive need to overeat highly palatable food after a period of abstinence, similar to cravings that arise during drug abstinence in chronic drug users (Pickering et al., 2009; Sharma et al., 2013). Dopamine has been implicated in mediating compulsive behavior and craving in both drugs of abuse and overeating in obesity (Baik, 2013; Salamone, 2003; Small et al., 2003; Volkow & Wise, 2005). Like drugs of abuse, palatable food activates the dopaminergic system and acts in a similarly reinforcing manner (Volkow et al., 2017), activating dopaminergic associated reward areas such as the nucleus accumbens (NAc), the ventral tegmental area, the hypothalamus, and the prefrontal cortex (Briggs et al., 2010). The dorsal striatum, though less studied than the NAc in appetitive motivation and addiction, has been associated with later, habitual stages of addiction (Everitt & Robbins, 2013; Volkow et al., 2006) and is believed to be a primary substrate mediating craving during abstinence that later leads to relapse (Contreras-Rodríguez et al., 2017; Volkow et al., 2006).

The neural substrates and mechanisms mediating the persistent craving that drives relapse in drug abstinence and dieting are only partially understood. Work in the last decade has demonstrated that drugs of abuse upregulate silent synapses, activating a substrate for circuit plasticity and remodeling that allows motivational circuits to become reorganized around behaviors associated with drug-taking (Dong, 2016; Graziane et al., 2016; X. Huang et al., 2015). Silent synapses contain functional NMDA but not AMPA receptors (Hanse et al., 2013). Due to the magnesium block on NMDA at resting potentials, without AMPARs to depolarize the postsynaptic region in response to glutamate, NMDA cannot pass current, and thus, the synapse remains ‘silent’ in response to glutamate release. Prevalent in early development, silent synapses act as a substrate for synapse selection and circuit development but are minimally expressed in adulthood (Atwood & Wojtowicz, 1999; X. Huang et al., 2015; J. Isaac, 2003; J. T. R. Isaac et al., 1995, p. 199). Drugs of abuse, such as cocaine, morphine, and chronic nicotine upregulate silent synapses in medium spiny neurons (MSNs) in a drug- and pathway-specific manner. Prior studies have identified two mechanisms by which silent synapses can be generated in adult animals: (i) the *de novo* generation of new synapses, typically enriched in NR2B-subunit containing NMDARs (Y. H. Huang et al., 2009), and (ii) the internalization of AMPA receptors from existing, established synapses (Graziane et al., 2016). Cocaine upregulates silent synapses in direct but not indirect pathway MSNs via *de novo* generation of new, NR2B enriched NMDA only synapses while opioids upregulate silent synapse in the indirect but not direct pathway via the removal of AMPARs from established synapses (Graziane et al., 2016; Y. H. Huang et al., 2009). Chronic nicotine increases silent synapses in the indirect pathway via *de novo* generation of NR2B-containing NMDA only synapses (Xia et al., 2017). Thus, different drugs induce alterations in silent synapses, and presumably circuit plasticity, in different target neuronal populations and by different mechanisms, though the functional significance of these different patterns is poorly understood.

Evidence suggests that the un-silencing of silent synapses following drug withdrawal facilitates incubation and craving during abstinence, suggesting the possibility that these silent synapses effectively encode drug-related associations that are then strengthened when unsilenced (Ma et al., 2014, 2016). Curiously, after normalization of silent synapses during abstinence, re-exposure to drug can induce a rapid return of increased silent synapses specifically in neurons previously encoding drug-related experience (Koya et al., 2012). That drugs of abuse can upregulate silent synapses demonstrates that this developmental mechanism mediating circuit plasticity can be reactivated in adulthood. Here we investigate whether HFD and associated dietary-induced obesity upregulates silent synapses in a manner similar to drugs of abuse in the dorsolateral striatum. As noted above, we select the DLS because of its role in mediating craving during abstinence (Contreras-Rodríguez et al., 2017; Volkow et al., 2006). Additionally, though the NAc is commonly studied, the DLS plays a critical role in motivating feeding behavior (Small et al., 2003; Sotak et al., 2005). Indeed, in dopamine-deficient mice that do not feed themselves and will die unless administered L-DOPA that facilitates feeding, genetic rescue of dopamine in the dorsolateral striatum but not the NAc can rescue feeding (Sotak et al., 2005), indicating that the DLS plays a critical role in feeding related appetitive motivation. In a dietary induced obesity paradigm, we fed mice HFD over a period of several weeks and then used *ex-vivo* patch-clamp electrophysiology to investigate alterations in silent synapses in both direct- and indirect-pathway MSNs. In contrast to cocaine and opioids, which upregulate silent synapses in only one of these pathways (Graziane et al., 2016; Y. H. Huang et al., 2009), we find that obesity and HFD upregulate silent synapses in both pathways, though the time course of both emergence of upregulation and normalization on withdrawal of HFD are prolonged compared to drugs of abuse.

## MATERIALS AND METHODS

### Animal

Adult mice (10-11 weeks at start of experiment) of both sexes were used for all experiments. To identify D1-expressing medium spiny neurons (MSNs), mice with a floxed tdTomato (JAX strain: #007914) were crossed with mice expressing cre-recombinase under the D1-promoter (B6.Cg-Tg(Drd1a-cre)262Gsat/Mmcd). Mice used in experiments were hemizygous for both the cre and floxed tdTomato alleles (tdTomato^wt/fl^; D1-cre^wt/cre^). To identify D2-expressing MSNs, we used mice hemizygous for a green fluorescent protein transgene under control of the drd2 promoter (JAX strain: #020631). All lines were on a C57Bl/6J background. Mice were group housed in a 12h light/dark cycle facility. All experiments were approved by Queens College, CUNY, Institutional Animal Care and Use Committee in accordance with the National Institutes of Health (NIH) Guidelines for the responsible use of animals in research.

### HFD Feeding

Under dietary induced obesity (DIO), mice received *ad libitum* access to high fat diet (HFD, 60% total calories from fat, Envigo, TD.06414,). HFD was administered for a minimum of 6 weeks prior to electrophysiology. In the time course experiments, mice received HFD for the periods indicated prior to electrophysiology. In the studies of renormalization after discontinuation of HFD, mice received 6 weeks of HFD before being switched to standard chow for HFD discontinuation.

### Slice preparation

Mice were anesthetized using euthasol (.05mL i.p, injection) and decapitated. The brain was removed rapidly from the cranial cavity and placed into ice-cold (4 °C) artificial cerebrospinal fluid (ACSF), in mM: 125 NaCl, 2.5 KCl, 2 CaCl_2_, 1 MgCl_2_, 1.25 NaH_2_PO_4_, 26 NaHCO_3_, 12.5 glucose, and 1 Na-ascorbate, and maintained at pH 7.4 by oxygenating with 95% O2/5% CO_2_. Coronal slices (300μm) containing dorsal striatum were obtained using VT1000 S vibratome (Leica Biosystems, Buffalo Grove, IL). Slices were transferred and incubated for at least 60 minutes at a holding temperature of 32-34 °C in ACSF and continuously bubbled with 95% O2/5% CO_2_. Slices were then maintained at room temperature (~27 °C) for the remainder of experiments.

### Electrophysiology

Single hemispheric coronal slices (300um) were transferred post-incubation into the recording chamber and perfused with oxygenated ACSF at a constant flow rate of 1.5-2mL/min. Temperature of ACSF was maintained at 30±1 °C using automatic temperature control (TC-324B Warner Instrument, Hamden, CT). Whole-cell patch clamp recordings from tdTomato labeled D1-expressing and EGFP-labeled D2-expressing MSNs were performed as previously described (Augustin et al., 2014). Recording pipettes with 4-8 MΩ resistance were filled with an internal solution containing, in mM: 120 CsMeSO_3_, 15 CsCl, 8 NaCl, 0.2 EGTA, 10 HEPES, 2 Mg-ATP, 0.3 Na-GTP, 10 TEA, 5 QX-314, adjusted to pH 7.3 with CsOH. Picrotoxin (50 μM) was added to ACSF in all experiments to block GABA_A_ receptor-mediated synaptic currents.

Bipolar twisted tungsten matrix microelectrode (FH-Co, Bowdoin, ME) with 500μm tip separation was placed in the striatum, dorsolateral to the recording electrode location. Experiments were performed to establish the percentage of silent synapses in recorded neurons using the minimal stimulation assay, in which failure rates of EPSCs at −70 and +40 mV were compared. In brief, small (<40 pA) EPSCs were evoked at −70 mV (single pulse every 5 seconds), then stimulation intensity was reduced until failures and successes could be clearly distinguished and a failure rate of approximately 50% was obtained. EPSCs are recorded at this stimulation intensity (i.e., produces 50% failure rate) across 50 trials at −70 mV before the holding potential is switched +40 mV and another 50 trials are recorded using the same stimulation intensity. Successes and failures were identified visually. The percentage of silent synapses is calculated comparing failures at −70mV to failures at +40mV using the formula: 1−Ln(Failure_−70 mV_) / Ln(Failure_+40 mV_). Mini excitatory post synaptic currents (mEPSCs) were recorded using Multiclamp 700B amplifier at a holding potential of −70mV. Miniature EPSCs were recorded (5-min samples, gap-free recording) with picrotoxin (50 μM) and tetrodotoxin (10 μM) applied in bath.

For experiments reported in Fig.3, NMDA subunit specific currents were isolated in Mg^2++^-free ACSF. Test pulses were applied every 20 seconds (plot created using 1 minute average of 3 pulses) at −70mV. Evoked responses were measured under 3 conditions: 1) Five-minute baseline in ACSF containing the AMPAR antagonist NBQX (5 μM) (0-5), 2) the addition of Ro25-6981 to selectively block NR2B (min 6-16), and 3) the addition of NVP-AAM077 (50 nM) to additionally block NR2A (min 17–26). NBQX was present throughout the entire recording.

### Spine morphology/Dil labeling

Mice were anesthetized by intraperitoneal injection of Euthasol and perfused transcardially with PBS and 1.5% paraformaldehyde (PFA) in PBS. Brains were removed and postfixed in 1.5% PFA for one hour. Coronal slices of 150 um were made using a vibratome and collected in PBS solution. Fluorescent DiI (1,1′-Dioctadecyl-3,3,3′,3′-tetramethylindocarbocyanine perchlorate (‘DiI’); DiIC18(3)) was introduced into the tissue by ballistic delivery using the PDS-1000/He System, using 1,100 psi rupture discs (Bio-Rad, California, #1652329). Once the dye was introduced into the tissue the tissue was stored in the dark at 4°C for 24 hours to allow for the dye to diffuse through the neuronal membranes. Dil was prepared as follows:

1.5 mg of DiI was dissolved in 50 ul of methylene chloride. The solution was then poured on 12.5 mg of tungsten particles and allowed to dry. The dye was then dissolved in 3 ml of deionized water and sonicated for 20 minutes. Afterwards one drop of the solution was added to the center of a microcarrier film and allowed to dry and then stored in the dark at 4°C.

### Immunohistochemistry and confocal imaging

Immunohistochemistry (IHC) was performed after DiI labeling to relabel D2-expressing MSNs. To label D2-expressing cells, one of either two IHCs were performed: (1) chicken anti-GFP to enhance the endogenous expression of GFP in D2 neurons or (2) rabbit anti-D2R to label D2 neurons. Both IHCs used an Alexa Fluor 488 secondary antibody as well: (1) goat anti-chicken Alexa Fluor 488 or (2) donkey anti-rabbit Alexa Fluor 488. Tissue was permeabilized in .01% Triton-X with PBS for 15 minutes, blocked in .01 Triton-X and a 10% normal serum with PBS for 30 minutes, then incubated in the primary antibody at 1:1000 in blocking solution for 2 hours. Tissue was then washed 3 times for 10 minutes each in PBS. The secondary antibody was applied at 1:1000 in blocking solution followed by 4 10-minute washes in PBS. A confocal microscope (Olympus Fv10i) was used to image neurons. Imaging was done with Alexa Fluor 488 (Excitation 495 nm; Emission 519 nm) and DiI (Excitation 549 nm; Emission 565) filters using 60x oil immersion lens.

### Dendritic Spine Analysis

Secondary and tertiary spine segments between 60-90 um were sampled as primary dendritic branches contain sparse amounts of spines. A maximum sampling of 3 branches were taken from each labeled cell. Spines were classified as follows: (1) stubby spines: less than 1 μM, no neck protrusion from dendrite (2) mushroom spines: between 1-2 μM, with neck and head (3) thin spines: length between 2-3 μM with neck and head (4) filopodia: neck greater than 3 μM, no head. Analysis was completed using Neurolucida. One-way ANOVAs were used to compare differences in spine density and type.

## RESULTS

### Silent synapses are upregulated in the dorsolateral striatum under chronic HFD

Mice were placed on a HFD for a minimum of 6 weeks before electrophysiology testing. In a preliminary study, we demonstrate significant weight gain at 6 weeks (Fig 1A, *F*_(1,26)_ =34.37, *p*<.01), consistent with prior reports in mice that show that 5-6 weeks of HFD induces significant weight gain and alters metabolic regulation (Krishna et al., 2016). Silent synapses were assessed using the minimal stimulation assay targeting the dorsolateral striatum. To identify direct and indirect pathway medium spiny neurons (MSNs), we used mice expressing red fluorescent protein in D1-expressing MSNs (tdTomato x D1-Cre) or a green fluorescent protein under the Drd2 promotor (D2-EGFP).

**Fig 1.**
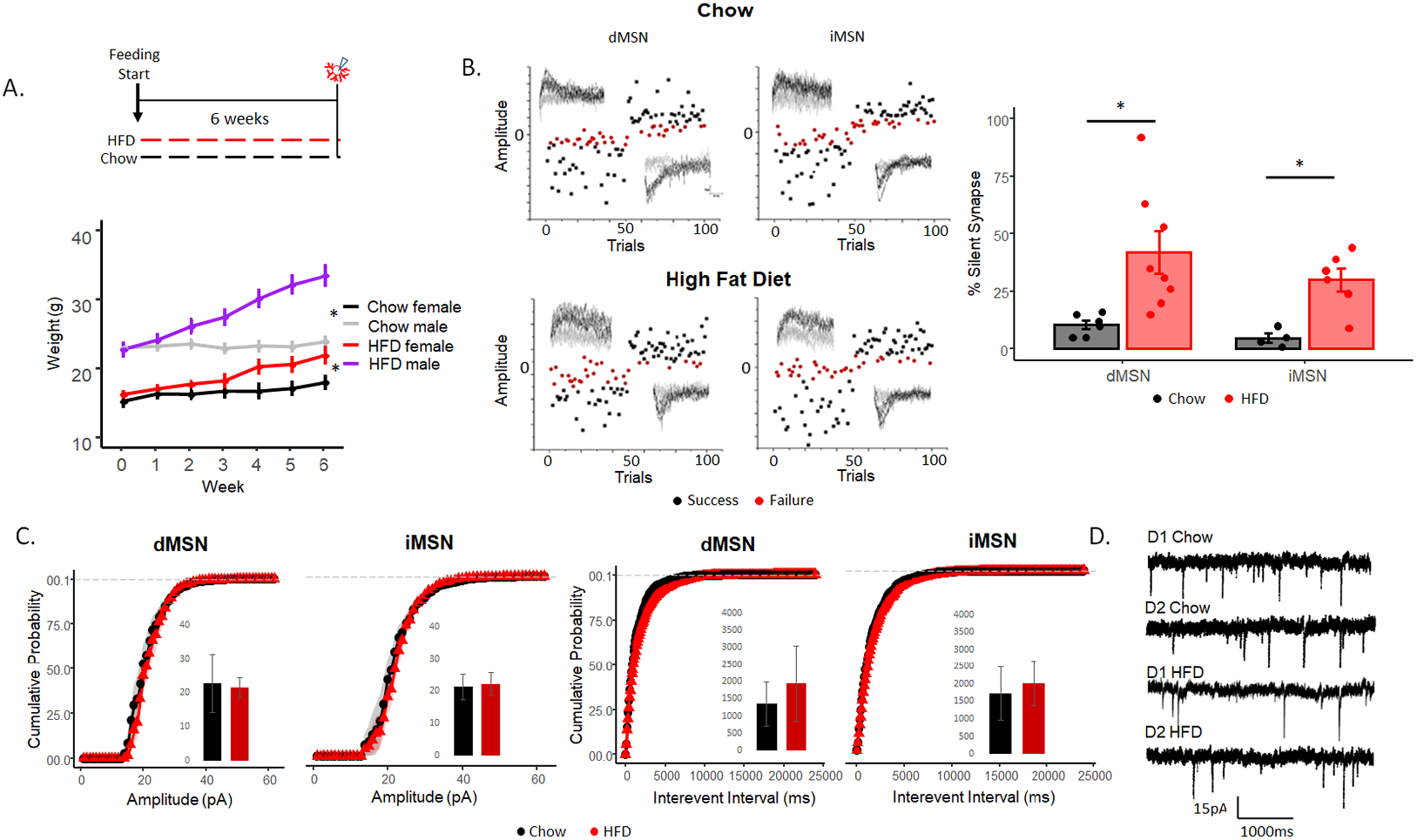
Silent synapses are upregulated in the dorsolateral striatum under chronic HFD. A) Feeding paradigm for chow and HFD mice (top) and weight changes over time (bottom). B) Minimal stimulation assay (left) and the percentage of silent synapses (right) of dMSNs and iMSNs in chow and HFD-fed mice. C) Cumulative probability of amplitude (pA) (left) and interevent interval (ms) (right) in dMSN and iMSNs. D) Example traces of gap-free mEPSCs with 50 μM picrotoxin and 10 μM TTX. Data are presented as mean ± SEM. *p< 0.05.

Silent synapses were increased in mice on HFD compared to controls (Fig. 1B, *F*_(1,22)_=13.84, *p*<.01) in both direct-(Fig. 1B, *F*_(1,12)_ = 6.89, *p*<.05) and indirect-(Fig. 1B, *F*_(1,8)_ = 15.02, *p*<.05) pathway MSNs with no sex differences observed (data not shown, *F*_(1,20)_ = .21, *p=*.64). To evaluate whether HFD induced generalized changes in glutamatergic synaptic transmission, we measured miniature EPSCs (mEPSCs) in gap-free recordings to assess pre- and postsynaptic changes in synaptic transmission of AMPA-containing (i.e., not silent) synapses at −70mV. HFD did not alter the frequency of miniature EPSCs in the direct (Fig 1D-E, *F*_(1,16)_=.873, p=.363) or indirect (*F*_(1,13)_=.443, p=.52) pathways, suggesting no changes in the presynaptic probability of release. Amplitude was not altered in either the direct (Fig 1D-E, *F*_(1,16)_= .26, p=.61) or indirect (*F*_(1,13)_= .06, p=.8,) pathway, suggesting no alteration in postsynaptic glutamate response among established AMPA-containing synapses. Decay of mEPSCS were assessed and no difference was observed between chow and HFD mice (direct: *t*(21)=0.41, *p*=.68, indirect: *t*(15)=0.79, *p*=.47). These data suggest that HFD increases NMDA-only silent synapses without altering presynaptic glutamate release or the strength of transmission in existing non-silent synapses.

### Upregulation of silent synapses under HFD is delayed compared to drugs of abuse

Cocaine and morphine induce upregulate silent synapses in MSNs within 5-7 days of once-daily injections and, upon drug discontinuation, this upregulation normalizes to baseline (5-15% silent synapses) within 7 days of the last injection (Graziane et al., 2016). We assessed the onset of silent synapse upregulation under HFD by assessing silent synapses after 1-, 2-, and 4 weeks of HFD. There was no significant difference between HFD fed mice and controls across weeks 1-4 of HFD exposure in either pathway (Fig. 2A, 1 week: *F*_(1,8)_=.073, *p*=.79, 2 weeks: *F*_(1,8)_=2.69, *p*=.14, 4 weeks: *F*_(1,8)_=.038, *p*=.85). These data suggest the underlying mechanism mediating the increase in silent synapses operates on a slower timescale, reflecting more long-term adaptations.

**Fig 2.**
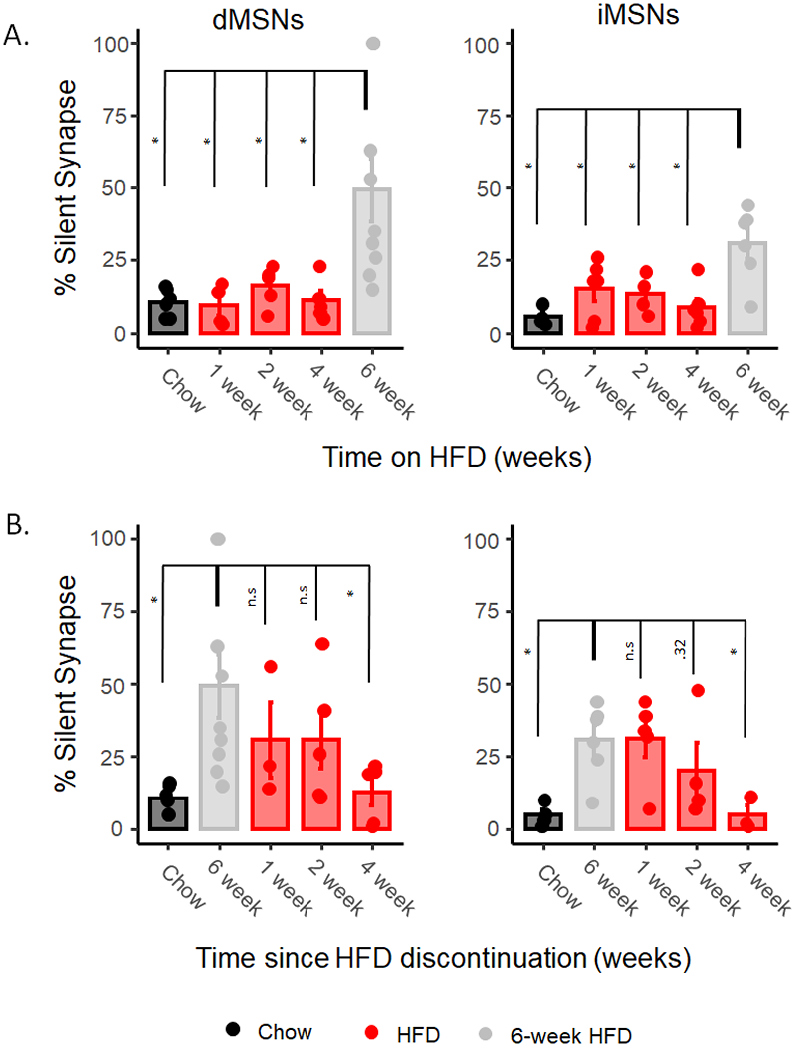
Upregulation of silent synapses under HFD is delayed compared to drugs of abuse. A) The onset of silent synapses after starting HFD feeding at 1 week, 2 weeks, and 4 weeks using minimal stimulation assay compared statistically to 6 weeks. B) the percentage of silent synapses after withdrawal from HFD at 1 week, 2 weeks, and 4 weeks as measured by minimal stimulation compared statistically to 6 weeks of HFD feeding. Data are presented as mean ± SEM. *p< 0.05.

We assessed silent synapses at 1-, 2-, and 4 weeks following withdrawal from HFD, observing a significant effect of time from withdrawal (dMSN: *F*_(4,23)_=3.39, *p*<.05; iMSN: *F*_(4,17)_=4.44, *p*<.05, Fig.2B). In post hoc comparisons examining the significance of differences between timepoints, neither the 1- or 2-week timepoint were significantly different from silent synapses in mice on HFD in either the direct (1 week: *F*_(1,10)_=.82, *p=*.38; 2 weeks: *F*_(1,12)_=1.25, *p=*.28) or indirect (1 week: *F*_(1,9)_=.004, *p=*.94; 2 weeks: *F*_(1,8)_=1.10, *p=*.32) pathway. By 4 weeks post withdrawal, silent synapses of mice withdrawn from HFD were not significantly different from chow-fed controls in both the direct or indirect pathway medium spiny neurons (Fig.2B, direct: *F*_(1,9)_=.239, *p*=.63; indirect: *F*_(1,5)_=.001, *p*=.98, respectively), indicating normalization to baseline similar to mice never exposed to HFD occurs approximately 4 weeks after discontinuation of HFD.

### HFD does not induce NR2B subunit enrichment in NMDARs

Increased silent synapses induced by drugs of abuse can arise from two different mechanisms. Cocaine induces increased silent synapses via putative *do novo* generation of new NR2B-subunit containing synapses, as suggested by enrichment in NR2B compared to control animals (Y. H. Huang et al., 2009; Xia et al., 2017). In contrast, morphine does not induce NR2B enrichment but increases silent synapses in the indirect pathway via internalization of AMPA receptors from existing, established synapses (Graziane et al., 2016).

We assessed the prevalence of NR2B-subunit containing NMDA receptors in HFD mice using Ro25-6981, an NMDA antagonist selective for NR2B-containing NMDA receptors. We isolated NMDA current by applying NBQX and picrotoxin to block AMPARs and GABA_A_ activity in a Mg^2++^-free bath ACSF and applied test pulses every 20s over 5 minutes to establish a baseline of NMDA EPSC response. We then applied Ro25-6981 to selectively block NR2B-containing NMDA currents and recorded for 10 minutes. If there is an enrichment of NR2B-containing NMDA receptors, as associated with *de novo* generation of new silent synapses, then Ro-25-6981 will incur a greater decrease in currents in HFD animals (Graziane et al., 2016; Y. H. Huang et al., 2009; Xia et al., 2017). We observed no difference in the effect of Ro-25-6981 in NMDA currents in either pathway (Fig 3B-C, calculated from the averages of the final 3 traces (1 min) before application of NVP; direct: F_1,8_=.21, *p=*.65; indirect: F_1,11_=.06, *p=*.81, respectively), suggesting no enrichment in NR2B-containing NMDA, arguing against increased silent synapses through *de novo* generation of new (silent) synapses as a primary mechanism of HFD induced increases in silent synapses. Finally, we applied NVP-AAM077, an NR2A blocker, to block NR2A-containing NMDA currents and again observed no significant difference between HFD mice and control mice in either pathway (Fig 3B, direct: F_1,8_=.12, *p=*.74; indirect: F_1,11_=.14, *p=*.72, respectively).

**Fig 3.**
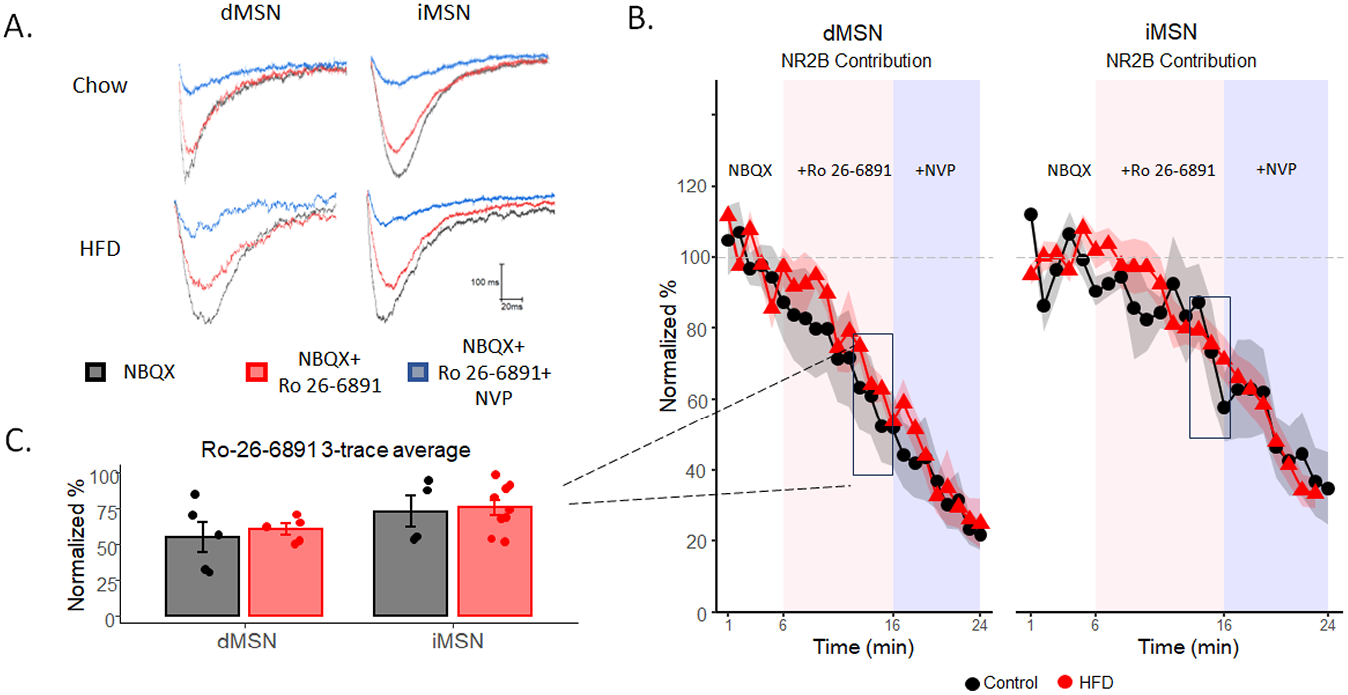
No increase in NR2B subunit prevalence is associated with upregulation of silent synapses. A) Example traces of stimulated EPSCs with the application of NBQX (black), NBQX and NR2B-subunit antagonist Ro 26-6891 (red), and NBQX, Ro 26-6891, and NR2A-subunit antagonist NVP (blue). B) Normalized percent change of NR2B contribution over time. C) The normalized percent at the end of Ro 26-6891 application (the average of 3 minutes) before the application of NVP. Data are presented as mean ± SEM. *p< 0.05.

### HFD reduces stubby dendritic spines in the indirect pathway

The plasticity of dendritic spines plays a role in shaping neural circuitry in an experience-dependent manner during both development and adulthood (Zito et al., 2009). Both exposure to HFD and withdrawal from drugs of abuse can alter spine density and morphology across brain region in a time- and region-dependent manner (Graziane et al., 2016; Saiyasit et al., 2020).

We examined dendritic spines in the dorsolateral striatum, comparing these in mice fed a HFD with chow-fed controls using diolistic imaging. DiO (red-dye) was incorporated into tissue of transgenic mice expressing green fluorescence under the Drd2 promotor (D2-GFP) to identify direct vs indirect pathway MSNs. Mice fed HFD for a minimum of six weeks showed a significant reduction in overall spine density (Fig. 4D left, *F*_1,128_=4.17, *p*<.043) in iMSN (*F*_1,58_=5.81, *p*<.05), but not dMSNs (*F*_1,70_=.18, *p=*.66). This density decrease in the indirect pathway is attributable specifically to the loss of stubby spines (Fig. 4D, *F*_1,58_=10.19, *p*<.05), with no significant change in spine density in either thin-long spines, mushroom spines, or filopodia in either pathway. Thin-long filopodia are highly labile and dynamic, whereas stubby and mushroom spines are more stable and mature. Changes in stubby spines can persist over longer periods of time, such as weeks compared to thin-long and filopodia. This decrease in stubby spines in iMSNs is consistent with the loss of mature, previously established synapses (Holtmaat et al., 2005).

**Fig 4.**
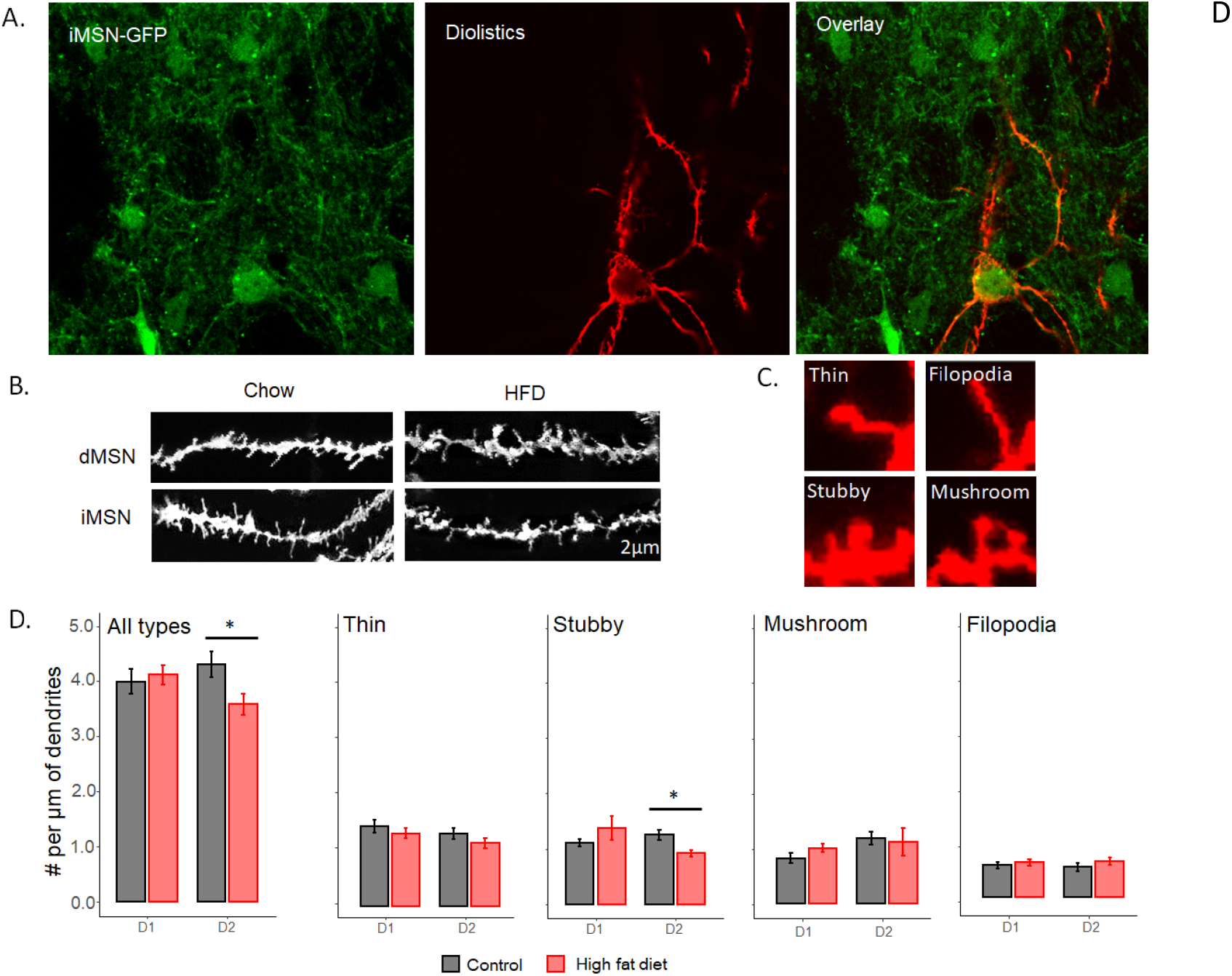
HFD reduces stubby dendritic spines in the indirect pathway. A) Representative images of iMSN-GFP neurons (left), DiI only (middle), and the overlap of iMSN-GFP and DiI labelling (right). B) Representative images of dendritic branch in dMSNs and iMSNs. C) Representative examples of different spine subtypes. D) Number of spines per μm of sampled dendritic branch including all spine subtypes (left) and broken down by individual spine subtype (right 4 graphs). Data are presented as mean ± SEM. *p< 0.05.

## DISCUSSION

Our study examined whether chronic HFD induces changes in silent synapse prevalence as observed with drugs of abuse. We find that HFD increases silent synapses in both direct and indirect pathway medium spiny neurons, in contrast to pathway selective effects of cocaine (D1R-expressing dMSNs) (Y. H. Huang et al., 2009), and morphine (D2R-expressing iMSNs) (Graziane et al., 2016). That silent synapses are increased in both pathways may indicate pervasive circuit remodeling associated with energy dense, highly palatable food in obesity. In drugs of abuse, modification of gene expression contributes to circuit remodeling (Bali & Kenny, 2019), with downregulation of the D2 receptor believed to be critical for drug mediated plasticity (Madhavan et al., 2013). Previous studies show that long-term consumption of HFD leads to alterations in *both* pathways, downregulating both D1 and D2 receptors in the NAc (Alsiö et al., 2010; Johnson & Kenny, 2010), which may facilitate circuit remodeling similarly to drugs of abuse. Our finding that HFD increases silent synapses is consistent with the parallel in these other mechanisms between neuroadaptations induced by drugs of abuse and HFD.

Our mEPSC examination of presynaptic and postsynaptic glutamatergic transmission in dMSNs and iMSNs after six weeks of HFD feeding showed no alteration in amplitude or interevent interval. As these studies were conducted at a holding potential of −70 mV, NMDA-only silent synapses with Mg^2++^ block at −70 mV were excluded, and the results reflect only the AMPAR current of non-silent synapses, suggesting that HFD did not induce a generalized change in glutamatergic transmission in either pathway. A previous study (Fritz et al., 2018) that examined the effect of HFD on mEPSCs in the DLS also found no difference in amplitude or frequency of mEPSCs between chow and HFD mice. That study observed a slower current decay in the HFD fed mice, though we did not observe an altered decay here, possibly due to the length of high fat feeding (6 weeks here vs 16 weeks in Fritz et al). The lack of changes in interevent interval and amplitude that we observe may also be region specific. Prior examinations of the effect of both cocaine (Corbit et al., 2014) and HFD (Fritz et al., 2018) on glutamate transmission in the DLS show no effect on interevent interval and amplitude, whereas both cocaine and HFD increase interevent interval in the NAc (Plaza-Briceño et al., 2023; Wang & Li, 2022).

We did not test AMPA/NMDA ratios as our focus is silent synapses which contain no AMPARs. Previous research has found that HFD can increase the AMPA/NMDA ratio in the DLS, though whether this is due to increased AMPARs or decreased NMDARs was not determined (Fritz et al., 2018). Notably, neither are consistent with an increase in silent synapses which would suggest an increase in NMDARs compared to AMPARs, either through *de novo* addition of NMDARs or through removal of AMPARs. However, research with morphine belies this simple assumption. Specifically, Saal et al (2003) demonstrate an increase in AMPA/NMDA ratio following morphine administration in the VTA. However, Graziane et al demonstrated that morphine administration generates silent synapses through internalization of AMPARs, which would be consistent with *decreased* AMPA/NMDA ratio. These studies, together, suggest that morphine, and analogously potentially HFD, may have multiple, complex effects on glutamate receptors, effects we were not able to isolate and characterize in this study.

Prior work with drugs of abuse have demonstrated distinct, pathway specific mechanisms mediating increased silent synapses: *de novo* creation of new synapses, typically enriched in NR2B (e.g., cocaine, direct pathway, (Y. H. Huang et al., 2009); chronic nicotine, indirect pathway, (Xia et al., 2017), or removal of AMPARs from established synapses (morphine, indirect pathway, (Graziane et al., 2016). As we observe an increase in silent synapses in *both* pathways, this provokes the question of which mechanism mediates this increase. As newly created silent synapses have been associated with NR2B enrichment, we tested for increased NR2B in HFD mice and found no difference in subunit composition in either pathway. Based on prior literature, this would suggest that HFD is not inducing *de novo* creation of new synapses like cocaine or nicotine. To demonstrate removal of AMPARs as the mechanism generating silent synapses following morphine administration, Graziane et al (2016) administered TAT-GluA2_3Y_, which blocks AMPAR internalization, just prior to each morphine administration and show that this prevents upregulation of silent synapses. In those studies, it was feasible to administer TAT-GluA2_3Y_ with each morphine injection. In the present study, mice were exposed to and ‘self-administered’ HFD 24 hours a day every day over a period of weeks, not providing a discrete administration with which to co-administer TAT-GluA2_3Y_. Even if feasible, were we to block AMPAR internalization over a period of several weeks, the effects would likely be pervasive and profound and not easily interpretable specifically with regard to silent synapses. Thus, our data suggest that HFD induced silent synapses are not mediated by de novo generation of NR2B enriched new synapses, which points to generation by removal and deprecation of existing synapses, though we cannot demonstrate that conclusively.

Silent synapse upregulation induced by HFD emerges more slowly compared to upregulation observed in response to drugs of abuse. Cocaine and morphine upregulate silent synapses between 5-7 days with daily administration (Graziane et al., 2016) and chronic nicotine exposure increases silent synapses in at most 21 days (Xia et al., 2017), possibly sooner. In the DLS, increases in silent synapses require greater than 4 weeks of high fat consumption. This delayed time course suggests that the changes in silent synapses may emerge secondary to slower adaptations induced by HFD; for example, emerging insulin resistance. Indeed, work by Plitzko et al (2001) demonstrates that insulin can regulate silent synapses in neocortical neurons during development. Similarly, the timecourse of normalization of silent synapses is delayed compared to drugs of abuse which occurs in approximately seven days for both cocaine and morphine (Graziane et al., 2016; Y. H. Huang et al., 2009). Here we observe continued upregulated silent synapses at 2 weeks after discontinuation of HFD that normalizes by 4 weeks. This suggests the normalization of upregulated silent synapses parallels a slower timecourse normalization of an underlying HFD induced adaptation. Withdrawal from cocaine recruits calcium permeable AMPA receptors to previously silent synapses, ‘unsilencing’ them (Shukla et al., 2017). In rats on a cafeteria diet, CP-AMPARs are increased following brief periods of withdrawal (Alonso-Caraballo et al., 2021), which could reflect a process of unsilencing silent synapses, though this effect was only seen in males and the authors did not examine silent synapses. While we did not determine the underlying mechanisms generating this more extended timecourse of the emergence and normalization of silent synapses, the contrasting timecourse observed between drugs of abuse and HFD as inducers of upregulated silent synapses could facilitate future investigations into regulatory mechanisms governing silent synapses in adulthood, still poorly understood.

Silent synapses have been associated with changes in spine density and morphology. Cocaine increases (Graziane et al., 2016) whereas morphine decreases spine density (Robinson et al., 2002) in the NAc. These changes in spine density are consistent with the *de novo* addition of new silent synapses generated by cocaine or the removal of AMPARs from established synapses--silencing them--induced by morphine. Prior studies have demonstrated varying effects of HFD on spine density in different brain regions. In one study, three week of HFD reduced spine density in the prefrontal cortex but not in the NAc (Dingess et al., 2017). In another study, HFD decreased spine density in the hippocampus (Hao et al., 2016). A ‘cafeteria’ diet decreased spine in the orbitofrontal cortex (Thompson et al., 2017). Here we observe pathway specific reductions in spine density in the dorsolateral striatum. Silent synapses have been suggested to be transient structures that either are selected and matured as non-silent synapses or, in the opposite direction precede the complete removal of a synapses (Hanse et al., 2013; X. Huang et al., 2015). Here we observe a selective loss of stubby spines, considered to reflect stable, mature synapses (Holtmaat et al., 2005). This is consistent with increased silent synapses reflecting an on-going process of degrading existing synapses, first through AMPAR removal to generate silent synapse, then to complete removal of the synapse. We did not, however, observe a difference in mEPSCS. Fewer total synapses on a cell, even with unchanged release probability, could reduce the total number of release events reflected in reduced mEPSC frequency, which we did not observe. However, Thompson et al (2017) also observed decreased dendritic spines concomitantly with no changes in mEPSCs, suggesting that loss of spines may not correspond to an observable change in glutamate transmission, at least as measured through spontaneous quantal release events.

Interestingly, the different timecourse we observe with HFD compared to drugs of abuse in altering silent synapse may be reflected in spine changes as well. Thin-type and filopodia spines are highly labile, while stubby spines are less dynamic and tend to change on a timescale of weeks, compared to minutes or hours. Drugs of abuse, which elicit silent synapses in days, primarily find alterations in thin-type and filopodia spines (Graziane et al., 2016), while we find no significant changes in these spine types under HFD. In contrast, HFD elicits silent synapses over weeks, not days, and we find alterations in spine types with a timecourse that, correspondingly, arises over weeks. This leaves open the question of whether this reflects substantially different mechanisms mediating these effects between HFD and drugs of abuse, or whether extended drug exposure might also induce alterations in stubby and mushroom spine types that tend to change over a longer rather than shorter timescale.

Overall, our findings indicate that chronic consumption of HFD can, similarly to drugs of abuse, induce an upregulation of silent synapses that could provide a mechanism and substrate for plasticity by which energy-dense, highly palatable foods reorganize neural circuits to generate compulsive, addiction-like behavior around eating. In drugs of abuse, long periods of withdrawal often cause an incubation of craving, in which the desire for that substance increases over an initial but extended period of abstinence (Pickens et al., 2011). This incubation has been linked to silent synapses (Lee et al., 2013). Incubation of food craving has also been observed in rodents (Grimm, 2020), with research suggesting that longer periods of abstinence may lead to significantly higher levels of food-seeking (Madangopal et al., 2022). Future work might examine the relationship between HFD-induced silent synapses and putative incubation of food craving caused by dieting, potentially a significant contributor to high rates of dieting failure and concomitant relapse to overeating and weight regain.

## FUNDING

This work was supported by the Whitehall Foundation (2016-12-24, JAB), the National Institute on Drug Abuse (DA052871, JAB) and a PSC-CUNY Award, jointly funded by The Professional Staff Congress and The City University of New York (JAB).

## Notes

### Summary of Updates

The manuscript has been revised in response to initial review. Revisions primarily focus on discussion.

## REFERENCES

Alonso-Caraballo, Y., Fetterly, T. L., Jorgensen, E. T., Nieto, A. M., Brown, T. E., & Ferrario, C. R. (2021). Sex specific effects of “junk-food” diet on calcium permeable AMPA receptors and silent synapses in the nucleus accumbens core. Neuropsychopharmacology, 46(3), 569–578. 10.1038/s41386-020-0781-1

Alsiö, J., Olszewski, P. K., Norbäck, A. H., Gunnarsson, Z. E. A., Levine, A. S., Pickering, C., & Schiöth, H. B. (2010). Dopamine D1 receptor gene expression decreases in the nucleus accumbens upon long-term exposure to palatable food and differs depending on diet-induced obesity phenotype in rats. Neuroscience, 171(3), 779–787. 10.1016/j.neuroscience.2010.09.046

Atwood, H. L., & Wojtowicz, J. M. (1999). Silent synapses in neural plasticity: Current evidence. Learning & Memory, 6(6), 542–571.

Augustin, S. M., Beeler, J. A., McGehee, D. S., & Zhuang, X. (2014). Cyclic AMP and Afferent Activity Govern Bidirectional Synaptic Plasticity in Striatopallidal Neurons. Journal of Neuroscience, 34(19), 6692–6699. 10.1523/JNEUROSCI.3906-13.2014

Baik, J.-H. (2013). Dopamine signaling in food addiction: Role of dopamine D2 receptors. BMB Reports, 46(11), 519–526. 10.5483/BMBRep.2013.46.11.207

Bali, P., & Kenny, P. J. (2019). Transcriptional mechanisms of drug addiction. Dialogues in Clinical Neuroscience, 21(4), 379–387. 10.31887/DCNS.2019.21.4/pkenny

Briggs, D. I., Enriori, P. J., Lemus, M. B., Cowley, M. A., & Andrews, Z. B. (2010). Diet-Induced Obesity Causes Ghrelin Resistance in Arcuate NPY/AgRP Neurons. Endocrinology, 151(10), 4745–4755. 10.1210/en.2010-0556

Contreras-Rodríguez, O., Martín-Pérez, C., Vilar-López, R., & Verdejo-Garcia, A. (2017). Ventral and Dorsal Striatum Networks in Obesity: Link to Food Craving and Weight Gain. Biological Psychiatry, 81(9), 789–796. 10.1016/j.biopsych.2015.11.020

Corbit, L. H., Chieng, B. C., & Balleine, B. W. (2014). Effects of Repeated Cocaine Exposure on Habit Learning and Reversal by N-Acetylcysteine. Neuropsychopharmacology, 39(8), 1893–1901. 10.1038/npp.2014.37

Dingess, P. M., Darling, R. A., Kurt Dolence, E., Culver, B. W., & Brown, T. E. (2017). Exposure to a diet high in fat attenuates dendritic spine density in the medial prefrontal cortex. Brain Structure and Function, 222(2), 1077–1085. 10.1007/s00429-016-1208-y

Dong, Y. (2016). Silent Synapse-Based Circuitry Remodeling in Drug Addiction. International Journal of Neuropsychopharmacology, 19(5), pyv136. 10.1093/ijnp/pyv136

Dulloo, A. G., Jacquet, J., & Girardier, L. (1997). Poststarvation hyperphagia and body fat overshooting in humans: A role for feedback signals from lean and fat tissues. The American Journal of Clinical Nutrition, 65(3), 717–723. 10.1093/ajcn/65.3.717

Everitt, B. J., & Robbins, T. W. (2013). From the ventral to the dorsal striatum: Devolving views of their roles in drug addiction. Neuroscience & Biobehavioral Reviews, 37(9), 1946–1954. 10.1016/j.neubiorev.2013.02.010

Flegal, K. M., Kruszon-Moran, D., Carroll, M. D., Fryar, C. D., & Ogden, C. L. (2016). Trends in Obesity Among Adults in the United States, 2005 to 2014. JAMA, 315(21), 2284. 10.1001/jama.2016.6458

Fritz, B. M., Muñoz, B., Yin, F., Bauchle, C., & Atwood, B. K. (2018). A High-fat, High-sugar ‘Western’ Diet Alters Dorsal Striatal Glutamate, Opioid, and Dopamine Transmission in Mice. Neuroscience, 372, 1–15. 10.1016/j.neuroscience.2017.12.036

Graziane, N. M., Sun, S., Wright, W. J., Jang, D., Liu, Z., Huang, Y. H., Nestler, E. J., Wang, Y. T., Schlüter, O. M., & Dong, Y. (2016). Opposing mechanisms mediate morphine- and cocaine-induced generation of silent synapses. Nature Neuroscience, 19(7), 915–925. 10.1038/nn.4313

Grimm, J. W. (2020). Incubation of food craving in rats: A review. Journal of the Experimental Analysis of Behavior, 113(1), 37–47. 10.1002/jeab.561

Hanse, E., Seth, H., & Riebe, I. (2013). AMPA-silent synapses in brain development and pathology. Nature Reviews Neuroscience, 14(12), 839–850. 10.1038/nrn3642

Hao, S., Dey, A., Yu, X., & Stranahan, A. M. (2016). Dietary obesity reversibly induces synaptic stripping by microglia and impairs hippocampal plasticity. Brain, Behavior, and Immunity, 51, 230–239. 10.1016/j.bbi.2015.08.023

Holtmaat, A. J. G. D., Trachtenberg, J. T., Wilbrecht, L., Shepherd, G. M., Zhang, X., Knott, G. W., & Svoboda, K. (2005). Transient and Persistent Dendritic Spines in the Neocortex In Vivo. Neuron, 45(2), 279–291. 10.1016/j.neuron.2005.01.003

Huang, X., Stodieck, S. K., Goetze, B., Cui, L., Wong, M. H., Wenzel, C., Hosang, L., Dong, Y., Löwel, S., & Schlüter, O. M. (2015). Progressive maturation of silent synapses governs the duration of a critical period. Proceedings of the National Academy of Sciences, 112(24), E3131–E3140. 10.1073/pnas.1506488112

Huang, Y. H., Lin, Y., Mu, P., Lee, B. R., Brown, T. E., Wayman, G., Marie, H., Liu, W., Yan, Z., Sorg, B. A., Schlüter, O. M., Zukin, R. S., & Dong, Y. (2009). In Vivo Cocaine Experience Generates Silent Synapses. Neuron, 63(1), 40–47. 10.1016/j.neuron.2009.06.007

Isaac, J. (2003). Postsynaptic silent synapses: Evidence and mechanisms. Neuropharmacology, 45(4), 450–460. 10.1016/S0028-3908(03)00229-6

Isaac, J. T. R., Nicoll, R. A., & Malenka, R. C. (1995). Evidence for silent synapses: Implications for the expression of LTP. Neuron, 15(2), 427–434. 10.1016/0896-6273(95)90046-2

Johnson, P. M., & Kenny, P. J. (2010). Dopamine D2 receptors in addiction-like reward dysfunction and compulsive eating in obese rats Supplementary Information. Nature Neuroscience, 13(5), 635– 641. 10.1038/nn.2519

Koya, E., Cruz, F. C., Ator, R., Golden, S. A., Hoffman, A. F., Lupica, C. R., & Hope, B. T. (2012). Silent synapses in selectively activated nucleus accumbens neurons following cocaine sensitization. Nature Neuroscience, 15(11), 1556–1562. 10.1038/nn.3232

Krishna, S., Lin, Z., Djani, D. M., Weber, M. T., Srivastava, L., & Filipov, N. M. (2016). Time-dependent behavioral, neurochemical, and metabolic dysregulation in female C57BL/6 mice caused by chronic high-fat diet intake. 13.

Lee, B. R., Ma, Y.-Y., Huang, Y. H., Wang, X., Otaka, M., Ishikawa, M., Neumann, P. A., Graziane, N. M., Brown, T. E., Suska, A., Guo, C., Lobo, M. K., Sesack, S. R., Wolf, M. E., Nestler, E. J., Shaham, Y., Schlüter, O. M., & Dong, Y. (2013). Maturation of silent synapses in amygdala-accumbens projection contributes to incubation of cocaine craving. Nature Neuroscience, 16(11), 1644– 1651. 10.1038/nn.3533

Ma, Y.-Y., Lee, B. R., Wang, X., Guo, C., Liu, L., Cui, R., Lan, Y., Balcita-Pedicino, J. J., Wolf, M. E., Sesack, S. R., Shaham, Y., Schlüter, O. M., Huang, Y. H., & Dong, Y. (2014). Bidirectional Modulation of Incubation of Cocaine Craving by Silent Synapse-Based Remodeling of Prefrontal Cortex to Accumbens Projections. Neuron, 83(6), 1453–1467. 10.1016/j.neuron.2014.08.023

Ma, Y.-Y., Wang, X., Huang, Y., Marie, H., Nestler, E. J., Schlüter, O. M., & Dong, Y. (2016). Re-silencing of silent synapses unmasks anti-relapse effects of environmental enrichment. Proceedings of the National Academy of Sciences, 113(18), 5089–5094. 10.1073/pnas.1524739113

Madangopal, R., Szelenyi, E. R., Nguyen, J., Brenner, M. B., Drake, O. R., Pham, D. Q., Shekara, A., Jin, M., Choong, J. J., Heins, C., Komer, L. E., Weber, S. J., Hope, B. T., Shaham, Y., & Golden, S. A. (2022). Incubation of palatable food craving is associated with brain-wide neuronal activation in mice. Proceedings of the National Academy of Sciences, 119(45), e2209382119. 10.1073/pnas.2209382119

Madhavan, A., Argilli, E., Bonci, A., & Whistler, J. L. (2013). Loss of D2 Dopamine Receptor Function Modulates Cocaine-Induced Glutamatergic Synaptic Potentiation in the Ventral Tegmental Area. Journal of Neuroscience, 33(30), 12329–12336. 10.1523/JNEUROSCI.0809-13.2013

Pickens, C. L., Airavaara, M., Theberge, F., Fanous, S., Hope, B. T., & Shaham, Y. (2011). Neurobiology of the incubation of drug craving. Trends in Neurosciences, 34(8), 411–420. 10.1016/j.tins.2011.06.001

Pickering, C., Alsiö, J., Hulting, A.-L., & Schiöth, H. B. (2009). Withdrawal from free-choice high-fat high-sugar diet induces craving only in obesity-prone animals. Psychopharmacology, 204(3), 431–443. 10.1007/s00213-009-1474-y

Plaza-Briceño, W., Velásquez, V. B., Silva-Olivares, F., Ceballo, K., Céspedes, R., Jorquera, G., Cruz, G., Martínez-Pinto, J., Bonansco, C., & Sotomayor-Zárate, R. (2023). Chronic Exposure to High Fat Diet Affects the Synaptic Transmission That Regulates the Dopamine Release in the Nucleus Accumbens of Adolescent Male Rats. International Journal of Molecular Sciences, 24(5), 4703. 10.3390/ijms24054703

Plitzko, D., Rumpel, S., & Gottmann, K. (2001). Insulin promotes functional induction of silent synapses in differentiating rat neocortical neurons. European Journal of Neuroscience, 14(8), 1412–1415.

Robinson, T. E., Gorny, G., Savage, V. R., & Kolb, B. (2002). Widespread but regionally specific effects of experimenter-versus self-administered morphine on dendritic spines in the nucleus accumbens, hippocampus, and neocortex of adult rats. Synapse, 46(4), 271–279. 10.1002/syn.10146

Saal, D., Dong, Y., Bonci, A., & Malenka, R. C. (2003). Drugs of abuse and stress trigger a common synaptic adaptation in dopamine neurons. Neuron, 37(4), 577–582. 10.1016/S0896-6273(03)00021-7

Saiyasit, N., Chunchai, T., Apaijai, N., Pratchayasakul, W., Sripetchwandee, J., Chattipakorn, N., & Chattipakorn, S. C. (2020). Chronic high-fat diet consumption induces an alteration in plasma/brain neurotensin signaling, metabolic disturbance, systemic inflammation/oxidative stress, brain apoptosis, and dendritic spine loss. Neuropeptides, 82, 102047. 10.1016/j.npep.2020.102047

Salamone, J. D. (2003). Nucleus Accumbens Dopamine and the Regulation of Effort in Food-Seeking Behavior: Implications for Studies of Natural Motivation, Psychiatry, and Drug Abuse. Journal of Pharmacology and Experimental Therapeutics, 305(1), 1–8. 10.1124/jpet.102.035063

Sharma, S., Fernandes, M. F., & Fulton, S. (2013). Adaptations in brain reward circuitry underlie palatable food cravings and anxiety induced by high-fat diet withdrawal. International Journal of Obesity, 37(9), 1183–1191. 10.1038/ijo.2012.197

Shukla, A., Beroun, A., Panopoulou, M., Neumann, P. A., Grant, S. G., Olive, M. F., Dong, Y., & Schlüter, O. M. (2017). Calcium-permeable AMPA receptors and silent synapses in cocaine-conditioned place preference. The EMBO Journal, 36(4), 458–474. 10.15252/embj.201695465

Small, D. M., Jones-Gotman, M., & Dagher, A. (2003). Feeding-induced dopamine release in dorsal striatum correlates with meal pleasantness ratings in healthy human volunteers. NeuroImage, 19(4), 1709–1715. 10.1016/S1053-8119(03)00253-2

Sotak, B. N., Hnasko, T. S., Robinson, S., Kremer, E. J., & Palmiter, R. D. (2005). Dysregulation of dopamine signaling in the dorsal striatum inhibits feeding. Brain Research, 1061(2), 88–96. 10.1016/j.brainres.2005.08.053

Speakman, J. R., Levitsky, D. A., Allison, D. B., Bray, M. S., de Castro, J. M., Clegg, D. J., Clapham, J. C., Dulloo, A. G., Gruer, L., Haw, S., Hebebrand, J., Hetherington, M. M., Higgs, S., Jebb, S. A., Loos, R. J. F., Luckman, S., Luke, A., Mohammed-Ali, V., O’Rahilly, S., … Westerterp-Plantenga, M. S. (2011). Set points, settling points and some alternative models: Theoretical options to understand how genes and environments combine to regulate body adiposity. Disease Models & Mechanisms, 4(6), 733–745. 10.1242/dmm.008698

Thompson, J. L., Drysdale, M., Baimel, C., Kaur, M., MacGowan, T., Pitman, K. A., & Borgland, S. L. (2017). Obesity-Induced Structural and Neuronal Plasticity in the Lateral Orbitofrontal Cortex. Neuropsychopharmacology, 42(7), 1480–1490. 10.1038/npp.2016.284

Volkow, N. D., Wang, G.-J., Telang, F., Fowler, J. S., Logan, J., Childress, A.-R., Jayne, M., Ma, Y., & Wong, C. (2006). Cocaine Cues and Dopamine in Dorsal Striatum: Mechanism of Craving in Cocaine Addiction. Journal of Neuroscience, 26(24), 6583–6588. 10.1523/JNEUROSCI.1544-06.2006

Volkow, N. D., & Wise, R. A. (2005). How can drug addiction help us understand obesity? Nature Neuroscience, 8(5), 555–560. 10.1038/nn1452

Volkow, N. D., Wise, R. A., & Baler, R. (2017). The dopamine motive system: Implications for drug and food addiction. Nature Reviews Neuroscience, 18(12), 741–752. 10.1038/nrn.2017.130

Wang, X., & Li, H. (2022). Chronic high-fat diet induces overeating and impairs synaptic transmission in feeding-related brain regions. Frontiers in Molecular Neuroscience, 15, 1019446. 10.3389/fnmol.2022.1019446

Xia, J., Meyers, A. M., & Beeler, J. A. (2017). Chronic Nicotine Alters Corticostriatal Plasticity in the Striatopallidal Pathway Mediated By NR2B-Containing Silent Synapses. Neuropsychopharmacology, 42(12), 2314–2324. 10.1038/npp.2017.87

Zito, K., Scheuss, V., Knott, G., Hill, T., & Svoboda, K. (2009). Rapid Functional Maturation of Nascent Dendritic Spines. Neuron, 61(2), 247–258. 10.1016/j.neuron.2008.10.054

